# *SeedExtractor*: an open-source GUI for seed image analysis

**DOI:** 10.1101/2020.06.28.176230

**Authors:** Feiyu Zhu, Puneet Paul, Waseem Hussain, Kyle Wallman, Balpreet K Dhatt, Jaspreet Sandhu, Larissa Irvin, Gota Morota, Hongfeng Yu, Harkamal Walia

## Abstract

Accurate measurement of seed size parameters is essential for both: breeding efforts□aimed at□enhancing yields and basic research□focused on discovering genetic components that regulate seed size. To address this need, we have developed an open-source graphical user interface (GUI) software, *SeedExtractor* that□determines seed size and shape (including area, perimeter, length, width, circularity, and centroid), and seed color with capability to process a large number of images in a time-efficient manner. In this context, our application takes ∼2 seconds for analyzing an image, i.e. significantly less compared to the other tools. As this software is open-source, it can be modified by users□to serve more specific needs. The adaptability of *SeedExtractor* was demonstrated by analyzing scanned seeds from multiple crops. We further validated the utility of this application by analyzing mature-rice seeds from 231 accessions in Rice Diversity Panel 1. The derived seed-size traits, such as seed length, width, were subjected to genome-wide association analysis. We identified well-known loci for regulating seed length (*GS3*) and width (*qSW5/GW5*) in rice, which demonstrated the accuracy of this application to extract seed phenotypes and accelerate trait discovery. In summary, we present a publicly available application that can be used to determine key yield-related traits in crops.

**HIGHLIGHT:** *SeedExtractor* is an open-source application designed to accurately measure seed size and seed color in a time-efficient manner for a wide variety of plant species.

## INTRODUCTION

Most of the plant-based food that we eat is either seed or seed-derived products. Thus, a large proportion of resources in crop improvement programs are invested towards better seeds. In this context, obtaining precise measurements of seed size and seed shape is critical to both: breeding programs aimed at enhancing crop yields, and facilitating fundamental research that is focused on discovering genetic components that serve to regulate seed size. Manual measurements of seed size provide evidence of restricted parameters such as length and width at a low resolution, which can be error-prone and time-consuming. Mechanized seed size measuring equipment is expensive, requires regular calibration, and often needs large amounts of seed to run through the system. In contrast, imaging-based automated platforms that are tailored to accurately measure seed parameters offer an efficient solution to minimize time constraints, seed amount issues, and circumvent manual errors. Moreover, high-throughput image analysis provides a powerful tool for trait discovery that facilities a more rapid input into downstream analysis such as genome-wide association studies (GWAS), that perform genetic mapping of yield-related traits.□

Qualitative assessment of the yield-related traits can also be important to ensure optimal nutritional values of seeds (Zhao *et al*., 2020). Within this framework, seed color can be associated with enhanced nutrition (Shao *et al*., 2011 and references therein). For instance, colored rice varieties carry antioxidant properties, which are known to decrease the risks involved with developing cardiovascular diseases (Ling *et al*., 2001). Similarly, pigmented maize seeds offer several beneficial effects on human health due to their antioxidant properties (Casas *et al*., 2014; Petroni *et al*., 2014). In addition to their medicinal properties, colored rice varieties hold cultural significance for certain regions and are consequentially valued in the respective local markets (Finocchiaro *et al*., 2007). Furthermore, the red pigmented wheat, which is resistant to pre-harvest sprouting, has been extensively targeted in wheat breeding programs (Groos *et al*., 2002).

Keeping in view the importance of seed size and color, several seed image analysis applications have been developed. For example, *SmartGrain* determines seed morphometrics such as area, perimeter, length, and width, as well as seed shape. However, it does not extract seed color information (Tanabata *et al*., 2012). On the other hand, *GrainScan* provides information with respect to seed size and color (Whan *et al*., 2014). Both the applications can be operated only on the windows platform. Although, these applications offer high levels of accuracy for analyzing seed images for size and shape determination, the adjustments that may be needed in setting the parameters are limited. For instance, *SmartGrain* only allows the user to determine the foreground and background colors, wherein *GrainScan* can only allow the user to set the size parameters. Moreover, processing a large number of images is time-consuming, and images with uneven illumination pose a challenge for precise measurements that may interfere with downstream analysis. These applications are not open-source and, therefore, cannot be further developed to improve based on user’s own needs.

To address the missing features in available seed image analysis software, we have developed a MATLAB based software tool –□*SeedExtractor*, an open-source graphical user interface (GUI) software tool that allows a user to conduct seed size analysis with precision. Based on the image processing libraries in MATLAB, our application is highly efficient, as it can process a large number of samples in a short period of time. The application allows the user to fine-tune the parameters for image processing and can handle a wide array of images. Most importantly, our application is open-source and MATLAB is available to most users through institutional license. This allows the user the ability to freely modify the application to suit more specific needs. As a test case to examine the value of this software, we screened mature seeds from 231 rice accessions corresponding to Rice diversity Panel 1 (RDP1) with different genetic (*indica, temperate japonica, tropical japonica, aus*, and *admixed*) and geographical backgrounds using□*SeedExtractor*. The derived seed-size related traits such as mature seed length and width were used to perform GWAS. Our□association mapping□confirmed the identity of known loci/genes regulating seed length (*GS3*) and width (*qSW5*) in rice, thus□validating the□accuracy of this application to□facilitate□genetic analysis and trait discovery.□

## MATERIALS AND METHOD

### □*SeedExtractor* workflow

*SeedExtractor*□is a MATLAB-based application, which makes it compatible with multiple operating systems. First, the MATLAB and *SeedExtractor* applications need to be installed. Then, the folders which contain the seed images (scanned or camera-based images) must be provided (Fig. 1). Next, the parameters, based on user’s requirement, is set and an individual image is tested to validate the optimal settings (Fig. 1). Sequentially, batch processing can be conducted to extract seed traits such as (1) area, (2) perimeter (3) major axis length (length), (4) minor axis length (width), (5) circularity, (7) seed number, (8) color intensity (different channels) and other digitally derived traits such as centroid. We have provided a step-by-step guide to use *SeedExtractor* (*SeedExtractor* Guide Document).

**Fig. 1.**
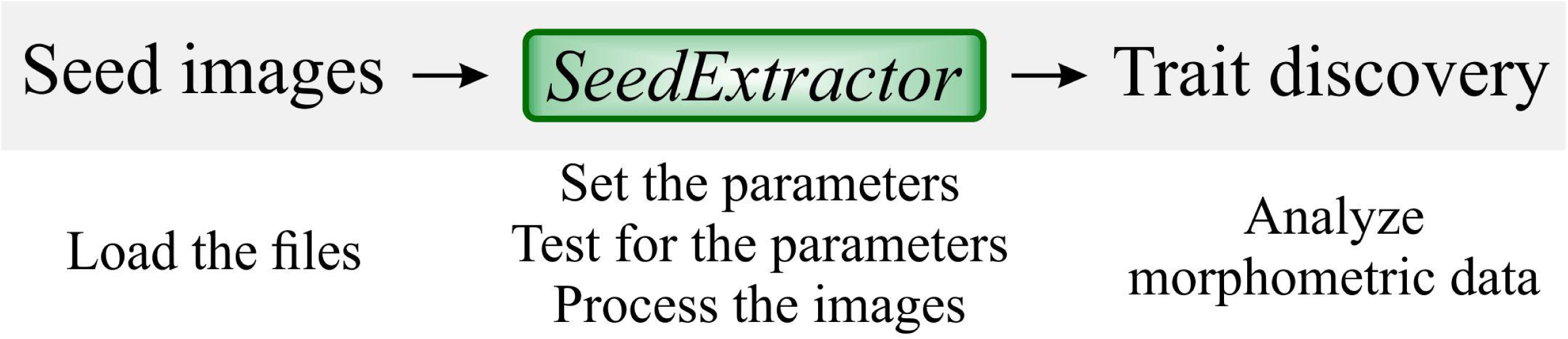
*SeedExtractor* workflow. Firstly, seed images are loaded, and the parameters are set. Testing of the parameters is performed to ensure optimal settings. Then, batch processing can be conducted to extract seed traits.

### Software implementation

#### Tool development

We have designed a GUI based on MATLAB, which provides the user the flexibility of setting unique parameters for processing seed images (Fig. 2).

**Fig.□2.**
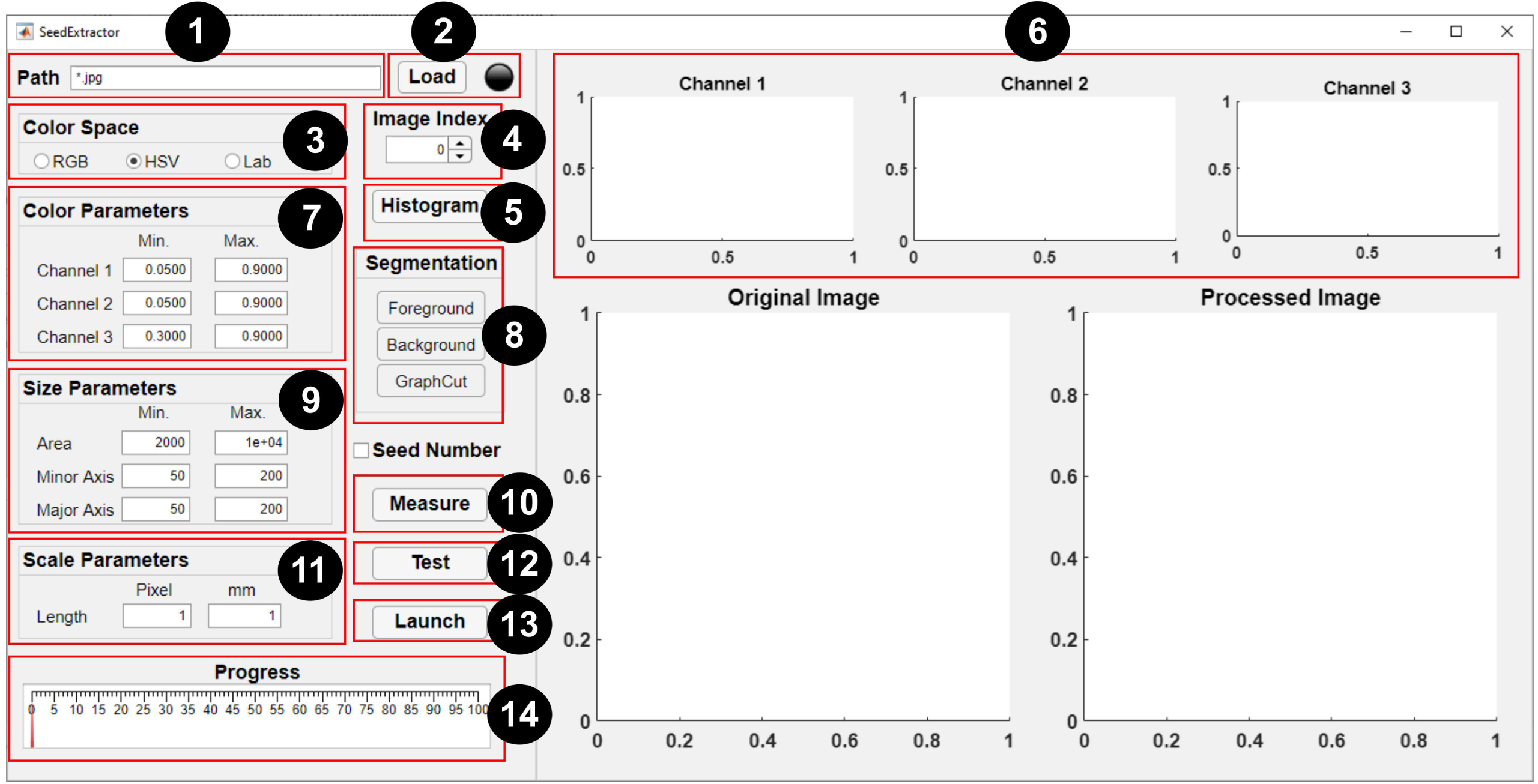
Graphical user interface of *SeedExtractor*.□The numbers denote a step-by-step guide on how to use the application: (1) path of the seed images is specified (* represents that all images in the particular folder need evaluation), (2) files are loaded automatically, (3) selection of color space should be made, (4) spinner can be used to change the current image (shown in the original image), (5) the user may select ‘histogram’ option (if applicable), (6) histograms representing distribution of colors in the three channels of the selected color space will be generated, (7) the range of histograms can be used to set the color parameters for the respective channel, (8) by selecting ‘foreground’ and ‘background’ – the user can scribble to define the color of the seed and background, respectively, and ‘graph cut’ will facilitate□segmentation of the seeds from the background, (9) minimum and maximum seed size parameters are defined (either default settings or manual corrections can be made) to filter out regions that are not seeds, (10) the user can ‘measure’□objects that have been used as a scale in the image and (11) define the scale measurement (in millimeters) that will aid in transforming the pixel length into metric units, (12) a test run should be performed□prior to batch processing in order to ensure that the parameter settings are optimized, (13) if the user has decided which parameters will be optimum,□batch□processing can be initiated, and (14) progress can be monitored via the progress bar.

#### Execution steps

A step by step guide is provided below to perform seed image analysis: (1) path specification, (2) file loading, (3) color space selection, (4) image selection, (5) histogram generation, (6) parameters setting, (7) graph cutting, (8) scale measurement, and (9) testing and processing.

##### Path specification

*SeedExtractor* is compatible with widely used image formats including *jpg*,□*png*, and□*tiff*. This□tool supports batch processing by loading all the images using a regular expression. For□example,□”FOLDER NAME*\\**.*jpg*”□loads□all the *jpg* images under the respective folder.

□□

##### File loading

Once□the□correct regular expression□has been typed□in ‘*Path*’ textbox,□the□’*Load*’ button can be clicked to load all the filenames into the application. The ‘*Light bulb*’□located□on the right side of the interface will turn red□while the filenames are being loaded.□Afterward, the□unprocessed□image will be shown in□’*Original Image*’ (Fig. 2). The spinner□can be used to change the index of the current image. The current image will be used for parameter setting and testing in later steps.

For accurate measurements, the ‘*Original Image*’ and ‘*Processed Image*’□can be zoomed in and out to check for any□discrepancy□between the original image and the processed image in the binary format. They can also be□panned□by holding the left-click button.□

□

##### Color space selection

The application□supports□three□different color□spaces: (1) red, green, and blue (*RGB*),□(2) hue, saturation, and value (*HSV*), (3) *Lab*. These three different choices of color spaces provide flexibility to the user in finding the optimal segmentation output. Once the color space is selected, the images will be processed in the respective color space for the next steps.

□

##### Histogram generation

The three histograms (*Channel 1, 2*, and *3*; Fig. 2) showing the distribution of colors in the three channels of the selected image (seed and the background) are generated. The meaning of the channels is dependent on the color space selected by the user. For example, if ‘*RGB*’ is chosen as a preferred color space, then histograms for ‘*Channel 1, 2*, and *3*’ refer to ‘*red, green*, and *blue*’. Similarly, ‘*hue, saturation*, and *value*’ for ‘*HSV*’, and ‘*l, a*, and *b*’ for ‘*Lab*’ color space. The distribution of colors in these three channels can be used as guide for setting the correct color ranges.□

□ □

##### Parameter setting

A set of default parameters are automatically loaded after launching the tool. Channel□ranges (minimum and maximum) are used to segment the seed regions from the background. Minimum and maximum seed size and shape parameters such as area, major and minor axis length, are used to filter out regions that are not seeds. However, the default parameters may not work for all the seed types or images. Thus, in this case, the user may need to set these parameters manually.

##### Graph cutting

To simplify the process of parameter setting, our application can also generate the parameters automatically based on ‘*user scribbles*’ to select the foreground and background. Then, using the ‘*GraphCut*’ algorithm (Kwatra *et al*., 2003), the foreground can be segmented from the background.

To select the foreground□(i.e., seed in this case), the user can click the ‘*foreground*’ button and scribble on□the seed□using a red mark□(Fig. 3a).□In cases where□the seed is too small, the user can zoom the image inward for scribbling.□Thereafter, ‘*Original Image*’ view can be restored.□To select the background, the user□can click the ‘*background*’ button□and scribble on the background using a green mark□(Fig. 3b).□

**Fig. 3.**
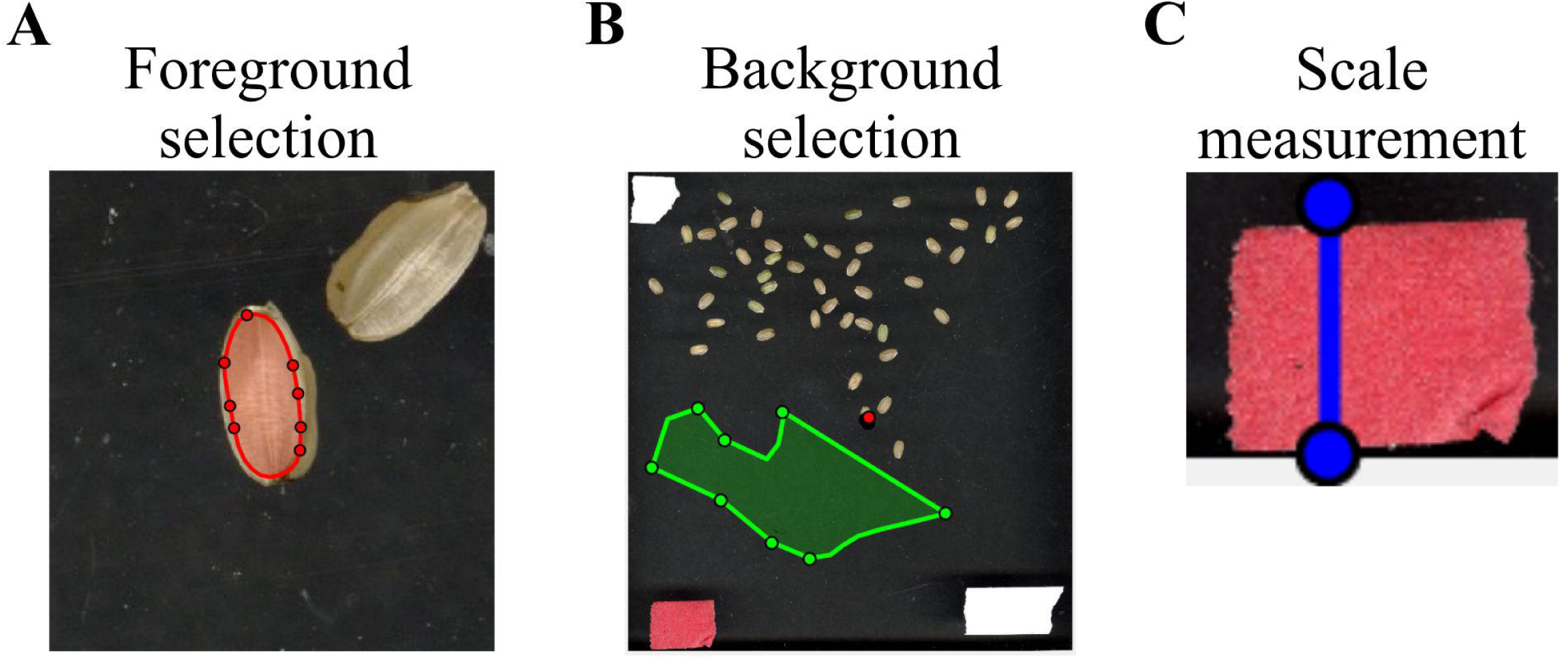
Selection of foreground, background, and scale measurements. By utilizing the function ‘*user scribbles*’, *SeedExtractor* can select foreground and background. (A) To select the foreground,□the user can click the ‘*foreground*’ button on the graphical user interface and scribble on□the seed□with a red mark. The image can be zoomed inward for the purpose of scribbling on smaller seeds. (B) For background selection, the user□can click the ‘*background*’ button□and scribble on the background with a green mark.□(C) For metric-scale measurements, the application allows the user to measure□objects that have been used as a scale in the image, which can then be used to transform the pixel length into millimeters. For this, a blue line can be drawn by clicking the ‘*Measure*’□button. When the line is drawn, the pixel length of the blue line will appear in the ‘*Length (pixel)*’ textbox. The user can type the corresponding length of the blue line in the ‘*Length (mm)*’ textbox. Then, the application automatically converts the selected values into metric units.

Once the foreground and background have been marked□or selected, the user can click the□’*GraphCut*’ button to□segment the seeds from the background. An image showing the mask of the foreground will be shown in□’*Processed Image*’ view. After selecting the ‘*GraphCut*’, the histograms corresponding only to the seed region will be displayed in□’*Channel 1, 2*, and *3*’□to guide the user in setting the color ranges.□Implementation of the ‘*GraphCut*’ function may take a few additional seconds. Supplementary Fig. S3 shows the histogram and parameter setting with and without ‘*GraphCut*’.

Due to a wide range of variation in seed size and color, it is difficult to automatically set optimal size and color ranges (for all of the color spaces). This tool provides the flexibility to the users to set these parameters manually based on the histograms. It is highly recommended that the user adjusts the parameters through testing. Nevertheless, the automatically generated parameters provide good initial values for the user to adjust accordingly.

##### Scale measurement

To obtain seed sizes in the metric system, the application allows the user to measure□objects that have been used as a scale in the image□(the tape in Fig. 3c). The known size of the scale can be used to transform the pixel length into millimeters (mm), thus presenting the extracted trait values into the metric system. For this, a blue line can be drawn by clicking the ‘*Measure*’□button. When the line is drawn, the pixel length of the blue line will appear in the ‘*Length (pixel)*’ textbox. The user can type the corresponding length of the blue line in the ‘*Length (mm)*’ textbox (Fig. 2). Then, the application automatically converts those values into metric units.

□

##### Testing□and processing

Once the user has set the parameters to□investigate how the parameters work, a test should be performed□prior to batch processing. To test the performance of the current parameters, the user can click the ‘*Test*’ button. An image showing the mask of the seeds will be□shown in□’*Processed Image*’ plot.□There is a checkbox□’*Seed Number*’, which is used to control whether□the seed regions in the processed image will be numbered or not. If the box is ticked, a series of□numbered yellow boxes will be drawn on the□lower right corners of the□individual seed□in the binary image.□

If the user has decided on the parameters to be used,□batch□processing can be initiated.□The processing will□begin□by clicking the□’*Launch*’□button. A series of traits will be□extracted□by the application, and the extracted traits will be□exported as□CSV□files.□The□’*Light bulb*’□will turn red during the processing of images and will turn green upon completion of the designated task. The ‘*Progress*’ gauge will show the progress of the image processing.

For each processed image, *SeedExtractor* will generate an output file that contains trait information of an individual seed in a particular image. Likewise, the mask of the seed regions from each image will be generated as a processed image. The indices of all the seed regions are marked in the processed image. In addition, the user can download combined file (TotalResult.csv) representing the average of particular trait for all the seeds per image from the MATLAB console.

□

## Algorithms

### Image Segmentation

The foreground with the seeds needs to be segmented from the background to process the image. We use the color thresholding technique to find the□seed regions.□We allow the user to segment the images in one□of□the□three color spaces,□RGB,□HSV, and□Lab. The default color space is□HSV, as we observed that HSV and Lab color spaces are better able to account for potentially uneven illuminations in the images. Range (minimum *C*_*i*_*min*_ and maximum *C*_*i*_*max*_) of the *i*th channel in the color parameter setting is□used to define the color ranges in the□selected□color□space.□More specifically, if□ *C*_1_, *C*_2_, and *C*_3_ □are the□three values of a pixel in the selected color space, a pixel satisfying the following inequalities will be identified as a seed pixel:

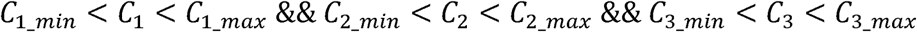

where && means the logic *and* operation. The processed image that is used as the mask of seed regions can be generated after color thresholding□in the selected color space (Bruce *et al*., 2000).□

□ The application detects each seed region in the binary format. The shape-related traits are□extracted from the binary or processed seed image and the colors are extracted from the original color image.□Currently, this application provides a series of traits such as seed number, area, perimeter, length, width, circularity, and centroid, as well as seed-color intensity.□□In this software, area *A*□is dictated by the number of pixels inside the region; perimeter *P*□is determined by the length of the boundary of the region.□Major (seed length) and□minor (seed width) axis lengths□are the lengths of the major and the minor axis□of the ellipse that has the same normalized second-central moments as the region. □Circularity□is□calculated as (4π*A*□)/*P*^2^ and can be used to evaluate how similar the region is to a circle. The□centroid□is the center of the seed region, which contains two values of coordinates.□Color intensities are the average intensity of Red, Green, and Blue channel intensity values for each seed region.□

### Performance Testing

To test the performance of *SeedExtractor*, we evaluated the time required to process: (case-I) images having a different number of seeds and (case-II) images at different levels of resolution (Supplementary Table S1 and S2, Supplementary Fig. S1). For this, mock seeds were computationally generated and increased from 1 seed to 100 seeds in a series of images (in case-I). In case-II, we used a fixed number of 10 seeds, and increased the level of resolution of each image from 50×50 to 1000×1000 pixels.

### Comparisons with other automated methods and manual measurements

First, we compared the time taken by *SeedExtractor* to analyze images (10 mature seed images from different rice) compared to other freely available applications such as *SmartGrain* and *GrainScan* (Supplementary Table S3). Next, we compared the accuracy of the seed morphometric measurements obtained by *SeedExtractor, SmartGrain*, and *GrainScan* to manual measurements using carbon fiber composite digital caliper (Resolution: 0.1 mm/0.01”, Accuracy: ±0.2 mm/0.01”, Power: 1.5 V; Fisherbrand). For this, we only considered seed length as it can be manually measured with relatively higher confidence levels than seed width. Raw values from manual and image-based measurements are provided in Supplementary Table S4. The comparison results will be detailed in the RESULTS and DISCUSSION section.

### Seed analyses from other plant species

To show adaptability of the application to measure seed images from other plant species, we analyzed images from rice, wheat, soybean, sorghum, common bean, and sunflower. These plant species represent a wide variation in the seed size.

### Rice Diversity Panel 1: a test case for *SeedExtractor* validation

Approximately 231 rice accessions from RDP1 (Liakat Ali *et al*., 2011; Zhao *et al*., 2011; Eizenga *et al*., 2014) were grown under optimal greenhouse conditions, 16 h light and 8 h dark at 28 ± 1°C and 23 ± 1°C, respectively, and a relative humidity of 55–60% (Dhatt *et al*., 2019). The harvested panicles were dried (30±1°C) for two weeks and mature seeds were dehusked using a Kett TR-250. The dehusked seeds were scanned using flatbed scanner – Epson Expression 12000 XL at 600 dpi resolution (Paul *et al*., 2020).□The seeds were spread out on a transparent plastic sheet placed on the glass of the scanner to avoid scratching. A□piece of tape at□0.5-inch□(12.7 mm)□width□was used for scaling.□

### Phenotypic analysis of morphometric measurements

*SeedExtractor* was used to obtain morphometric measurements on mature seed size. The various morphometric measurements derived from the scanned seed images were checked for normality and outliers were removed. The mature seed size data (length and width) was analyzed, and adjusted means for each accession across the replications were obtained with the following statistical model:

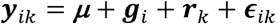

where ***y***_*ik*_ refers to the performance of the *i*th accession in the *k*th replication, ***μ*** is the intercept, ***g***_*i*_ is the effect of the *i*th accession, ***r***_*k*_ is the effect of *k*th replication, and ***ϵ***_*k*_ is the residual error associated with the observation ***y***_*ik*_. R statistical environment was used for the analysis (R Core Team, 2019).

### Genome wide association study (GWAS)

Adjusted means of various seed morphometric were used for GWAS analysis. GWAS was performed in rrBLUP R package (Endelman, 2011) using a high-density rice array (HDRA) of a 700k single nucleotide polymorphism (SNP) marker dataset (McCouch *et al*., 2016) with a total of 411,066 SNPs high quality SNPs retained after filtering out the missing data (< 20%) and minor allele frequency (< 5%). Following single marker linear mixed model was used for GWAS:

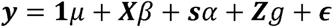

where **y** is a vector of observations, *µ* is the overall mean, **X** is the design matrix for fixed effects, ***β*** is a vector of principle components accounting for population structure, ***s*** is a vector reflecting the number of alleles (0,2) of each genotype at particular SNP locus, ***α*** is the effect of the SNP, ***Z*** is the design matrix for random effects, 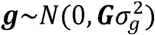, is the vector of random effects accounting for relatedness, **G** is the genomic relationship matrix of the genotypes, 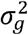 is the genetic variance, and ***ϵ*** is the vector of residuals. Manhattan plots were plotted using the qqman R package (Turner, 2014). To declare the genome-wide significance of SNP markers, we used a threshold level of *P* < 3.3 × 10^−6^ *p* or -log_10_(*P*) > 5.4 (Bai *et al*., 2016).

## RESULTS AND DISCUSSION

### Performance Test

We evaluated the performance of *SeedExtractor* with respect to the time required to process images. For this, we evaluated two cases: images having different numbers of seeds and images at different levels of resolution. In the first case, we used an incremental range (from 1 to 100) of seeds in a series of images (Supplementary Table S1, Supplementary Fig. S1A). We observed that the number of seeds does not affect the performance, as the time taken to process an image with 1 seed is similar to that of an image with 100 seeds (Fig. 4a). Secondly, we used a fixed number of seeds and increased the resolution of each consecutive image incrementally (Supplementary Table S2, Supplementary Fig. S1B). We detected that the performance of the application is slows gradually with increase in resolution as expected (Fig. 4b).

**Fig. 4.**
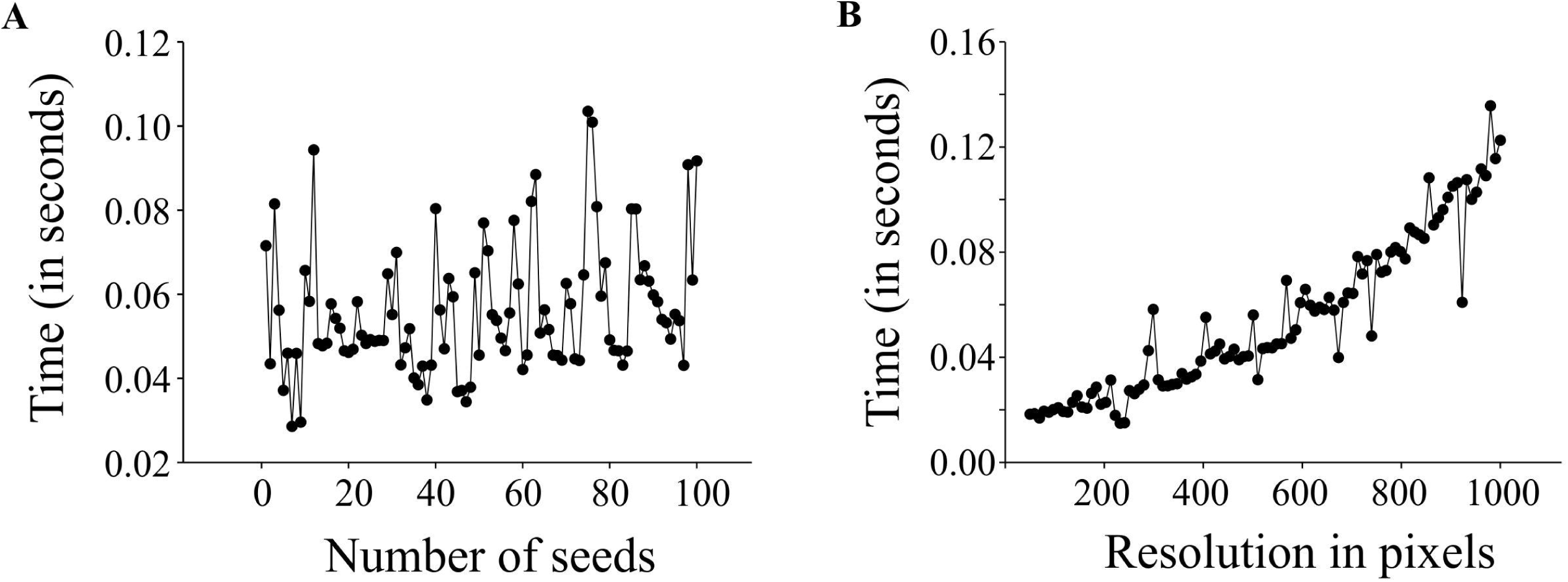
Performance testing of *SeedExtractor*. Plot showing the time taken to process images having different number of seeds (A), and images having different resolution levels (B).

### *SeedExtractor* versus other automated software and manual measurements

Next, we investigated the efficiency of *SeedExtractor* with respect to the time needed to analyze images relative to other automated software tools such as *SmartGrain* and *GrainScan*. Remarkably, the *SeedExtractor* takes ∼21 seconds for analyzing 10 images i.e., 30 times and 6 times more efficient than *SmartGrain* and *GrainScan*, respectively (Supplementary Table S3). Then, we correlated manual measurements with the analysis performed using each of the three automated softwares (*SeedExtractor, SmartGrain*, and *GrainScan*). Although manual measurements itself are prone to errors, we considered only seed lengths for the correlation because it can be measured with relatively higher confidence levels than seed width. Consequently, *SeedExtractor* showed the least deviation from manual measurements, as we detected correlation of 0.93 for *SeedExtractor*, 0.84 for *GrainScan*, and 0.92 for *SmartGrain* with manually measured seed length (Supplementary Table S4). Furthermore, we checked the correlation between the morphometric measurements obtained from the *SeedExtractor* and the other two softwares (Table 1, Supplementary Table S5). We detected a significantly high correlation (> 0.97) between the analyses conducted by *SeedExtractor* and *SmartGrain* (Table 1, Supplementary Table S5). Contrarily, the correlation between *GrainScan* and *SmartGrain* or *SeedExtractor* was relatively low (< 0.81; Table 1, Supplementary Table S5). Thus, *SeedExtractor* serves in a time-efficient and reliable manner to analyze seed size parameters.

**Table 1:** Correlation of the three automated applications for determining different seed size parameters.

| **Trait** | ***GrainScan***<br>**and**<br>***SmartGrain*** | ***GrainScan***<br>**and**<br>***SeedExtractor*** | ***SmartGrain***<br>**and**<br>***SeedExtractor*** |
| --- | --- | --- | --- |
| Area | 0.273 | 0.229 | 0.986 |
| Perimeter | 0.313 | 0.346 | 0.979 |
| Length | 0.549 | 0.518 | 0.995 |
| Width | 0.814 | 0.808 | 0.994 |
| Axis Ratio | NA | NA | 0.998 |
| Circularity | NA | NA | 0.923 |

For comparisons, mature seeds from one rice accession was considered. NA: not applicable

### Seed image analysis from other species

In addition to rice, seed measurements from other plant species representing a wide variation with respect to seed size were evaluated using *SeedExtractor*. For this, mature seeds from wheat, sorghum, common bean, and sunflower, were also carried out using *SeedExtractor*. After establishing the optimal parameters (Supplementary Fig. S2), *SeedExtractor* precisely segmented the mature seeds form the different plant species (Fig. 5). The successful and consistent derivation of the seed morphometrics from multiple plant species demonstrates the adaptability and utility of the application (Supplementary Table S6).

**Fig. 5.**
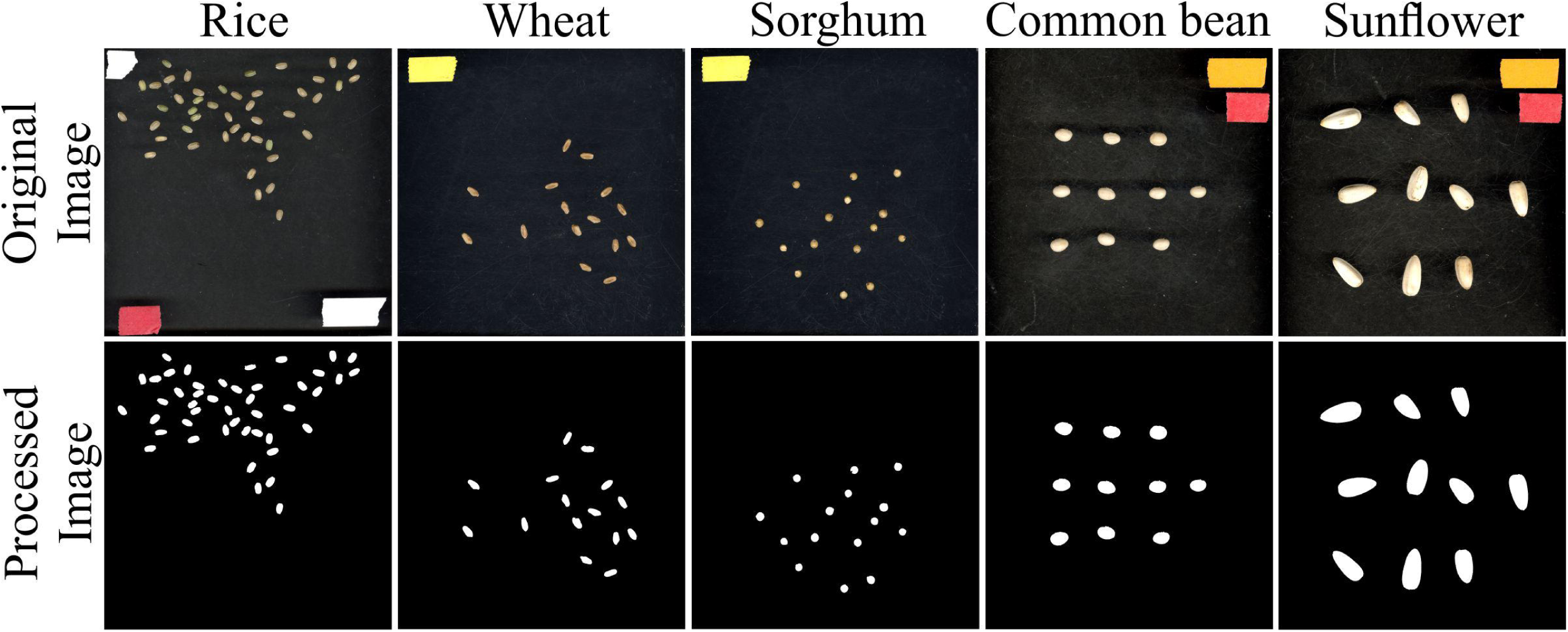
Seed analysis of different plant species. Mature seed images (original image) corresponding to rice, wheat, sorghum, common bean, and sunflower were evaluated using *SeedExtractor*. Processed image shows the segmented image pertaining to their respective plant species. Different color tapes in the original image were used for scaling purposes.

### Validation of *SeedExtractor* derived morphometric data

To validate the seed related traits derived from *SeedExtractor*, we screened 231 rice accessions corresponding to RDP1 (Supplementary Table S7). The mature seed length and width, which showed a normal distribution, were used for GWAS (Supplementary Fig. S4). Consequently, we identified 13 significant SNPs associated with seed length and 8 with seed width under control (Fig. 6, Supplementary Table S8). Remarkably, the lead SNP on chromosome 3 (SNP3.16732086; -log_10_*P* = 13.95) that affects mature seed length, corresponded to *GS3*, a known regulator of seed size (Fan *et al*., 2006). This known regulation of *GS3* was explanatory for 13.24% of phenotypic variation (Figure 6, Supplementary Table S8). *GS3* encodes a subunit of G-protein complex. Different alleles of *GS3* have been discussed to promote either longer (null alleles; Fan *et al*., 2006; Takano-Kai *et al*., 2009) or shorter seeds (gain-of-function allele; Mao *et al*., 2010). The other two significant SNPs for grain length were detected on chromosome 4 (SNP4.4655556; -log_10_*P* = 5.66) and 6 (SNP6.1112028; -log_10_*P* = 5.99), which encompasses *deformed interior floral organ 1* and an expressed protein, respectively.

**Fig. 6.**
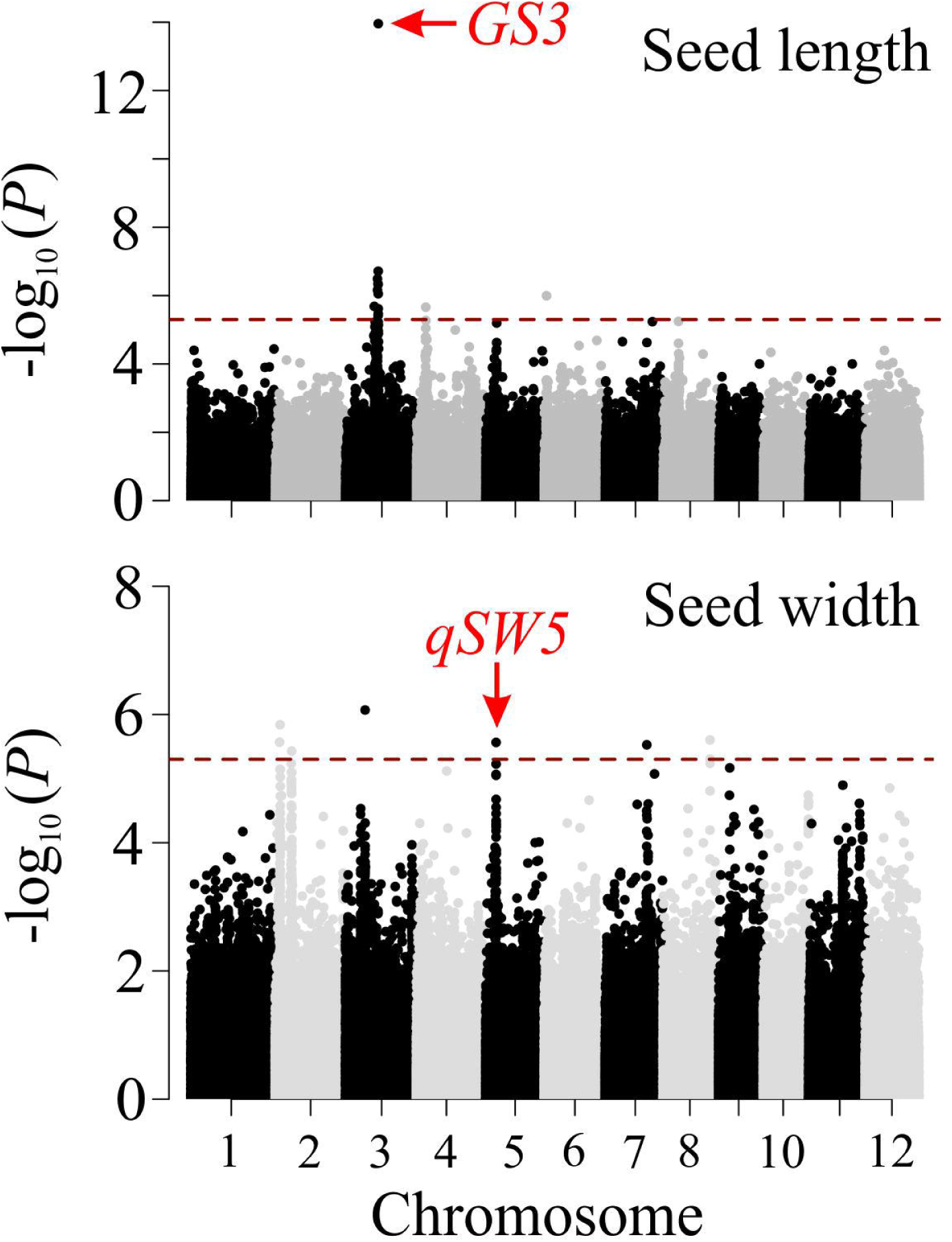
Manhattan plots of genome-wide association analysis for mature grain length (upper panel) and width (lower panel). The red dashed horizontal line indicates cut-off of significance threshold (*P* < 3.3 × 10^−6^ or -log_10_(*p*) > 5.4) level. Previously known major seed length (*GS3*) and width (*qSW5*) regulators are highlighted with a red arrow.

Furthermore, we identified several SNPs for seed width (Supplementary Table S8). For instance, the lead SNP on chromosome 2 (SNP2.2487459; -log_10_*P* = 6.07) co-localizes with an expressed protein (*Os02g05199*), and chromosome 3 (SNP3.10130641; -log_10_*P* = 5.84) is localized in the intergenic sequence between *Os03g18130* and *Os03g18140* (Figure 6, Supplementary Table S8). Interestingly, the significant SNP on chromosome 5 (SNP5.5348012; - log_10_*P* = 5.56; Figure 6, Supplementary Table S8) corresponded to a known regulator for seed width, *qSW5/GW5* (Weng *et al*., 2008; Duan *et al*., 2017; Liu *et al*., 2017; Kumar *et al*., 2019). This SNP explained phenotypic variation of 4.4%, which is in line with the previous studies (Huang *et al*., 2010; Zhao *et al*., 2011). The detection of the known seed size regulators, and the novel loci from the association mapping of the morphometric data, obtained by *SeedExtractor*, substantiates the power of the application to facilitate trait discovery. Collectively, these results validate the robustness of *SeedExtractor’s* ability to analyze seed size and seed shape parameters that can be used in downstream genetic analysis for trait discovery.

## CONCLUSION

This open-source cross-platform application provides a powerful tool to analyze seed images from a wide variety of plant species in a time-efficient manner. The accuracy of the tool is demonstrated by GWAS that identified the known regulators of seed length and width in rice. The versatility of this tool can extend beyond flatbed-scanned images, as it can also evaluate images taken by other cameras. In the future, this tool can be further developed to estimate other yield related parameters such as opaqueness or chalkiness in rice, which account for significant yield losses in global rice production.

## Supporting information

SeedExtractor Guide Document

## AUTHOR CONTRIBUTIONS

HW and HY supervised the project. PP lead the study. PP, BKD, JS, LI, and KW performed experiment on Rice Diversity Panel 1. PP scanned the seeds and performed manual measurements. FZ designed and developed the application. WH and GM performed analysis on the phenotypic data and genome-wide association mapping. PP and HW performed candidate gene analysis. PP and FZ wrote the manuscript. All authors read and approve the manuscript.

## FUNDING

This work was supported by National Science Foundation Award # 1736192 to HW, GM, and HY.

## ACKNOWLEDGEMENTS

We would like to thank Manny Saluja and Scott Sattler for providing sorghum seeds, Carlos Urrea for common bean seeds, Yavuz Delan and Ismail Dweikat for sunflower seeds.

## SOFTWARE AVAILABILITY

MATLAB can be downloaded from https://www.mathworks.com/products/matlab.html and *SeedExtractor* can be downloaded using the following link: https://cse.unl.edu/~fzhu/SeedExtractor.zip. We have provided detailed step-by-step guide for using *SeedExtractor* (*SeedExtractor* Guide Document).

## SUPPLEMENTARY MATERIAL

Supplementary Table S1: Performance testing with incremental seed numbers.

Supplementary Table S2: Performance testing with incremental image resolution.

Supplementary Table S3: Evaluation of efficiency with respect to time.

Supplementary Table S4: Manual and automated seed length measurements.

Supplementary Table S5: Morphometric analysis using the automated applications.

Supplementary Table S6: *SeedExtractor* based analysis of seed images from different plants.

Supplementary Table S7: Rice accessions used for genome wide association study.

Supplementary Table S8: Significant SNPs associated with seed length and width.

Supplementary Fig. S1: Images used for performance testing.

Supplementary Fig. S2: Parameters used to evaluate images from multiple plant species.

Supplementary Fig. S3: Graph-cutting.

Supplementary Fig. S4: Phenotypic distribution of mature seed length and width.

## Notes

### Competing Interest Statement

The authors have declared no competing interest.

