## Supplementary figures and images for "*SeedExtractor*: an open-source GUI for seed image analysis"

### Fig. S1

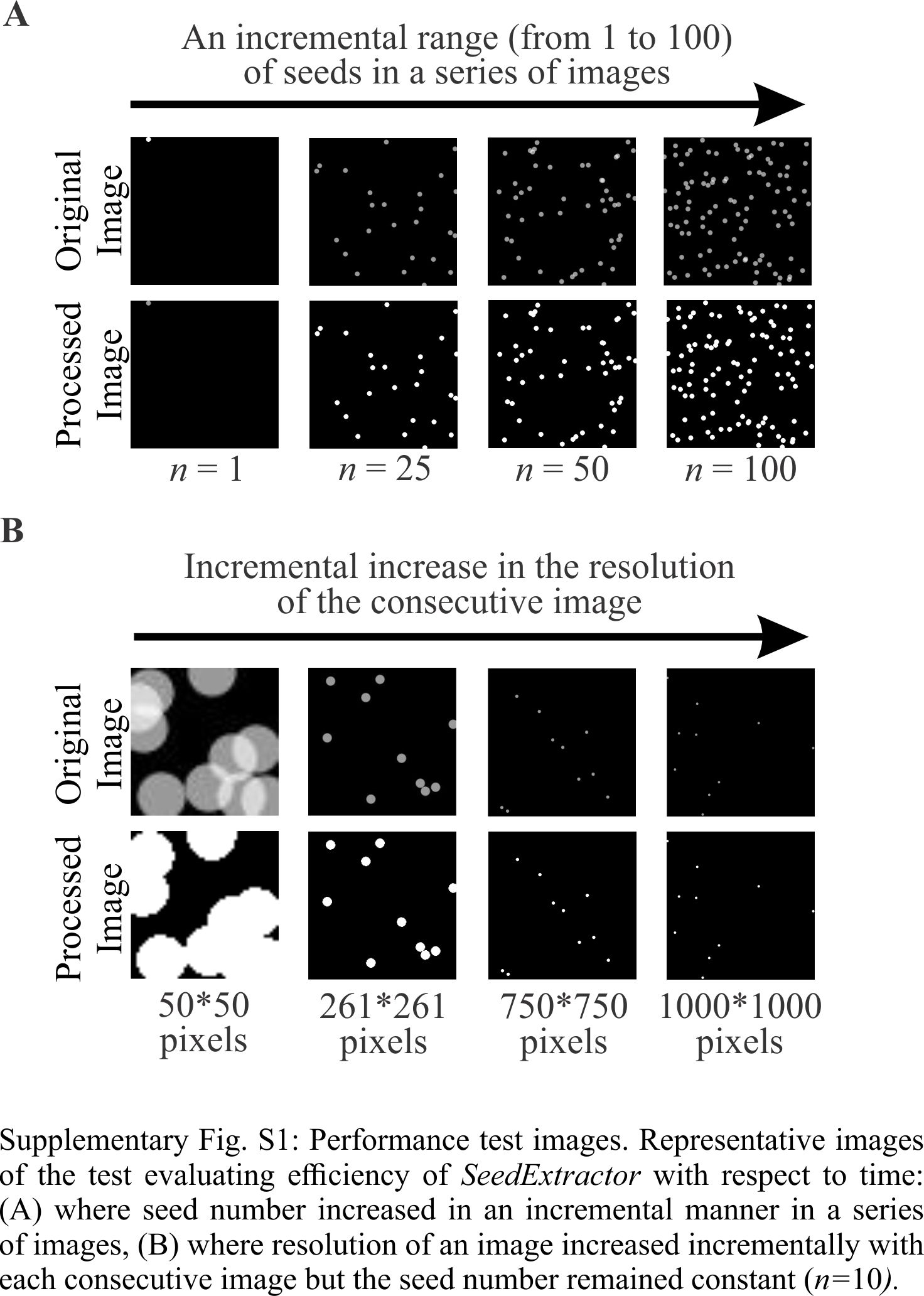

### Fig. S2

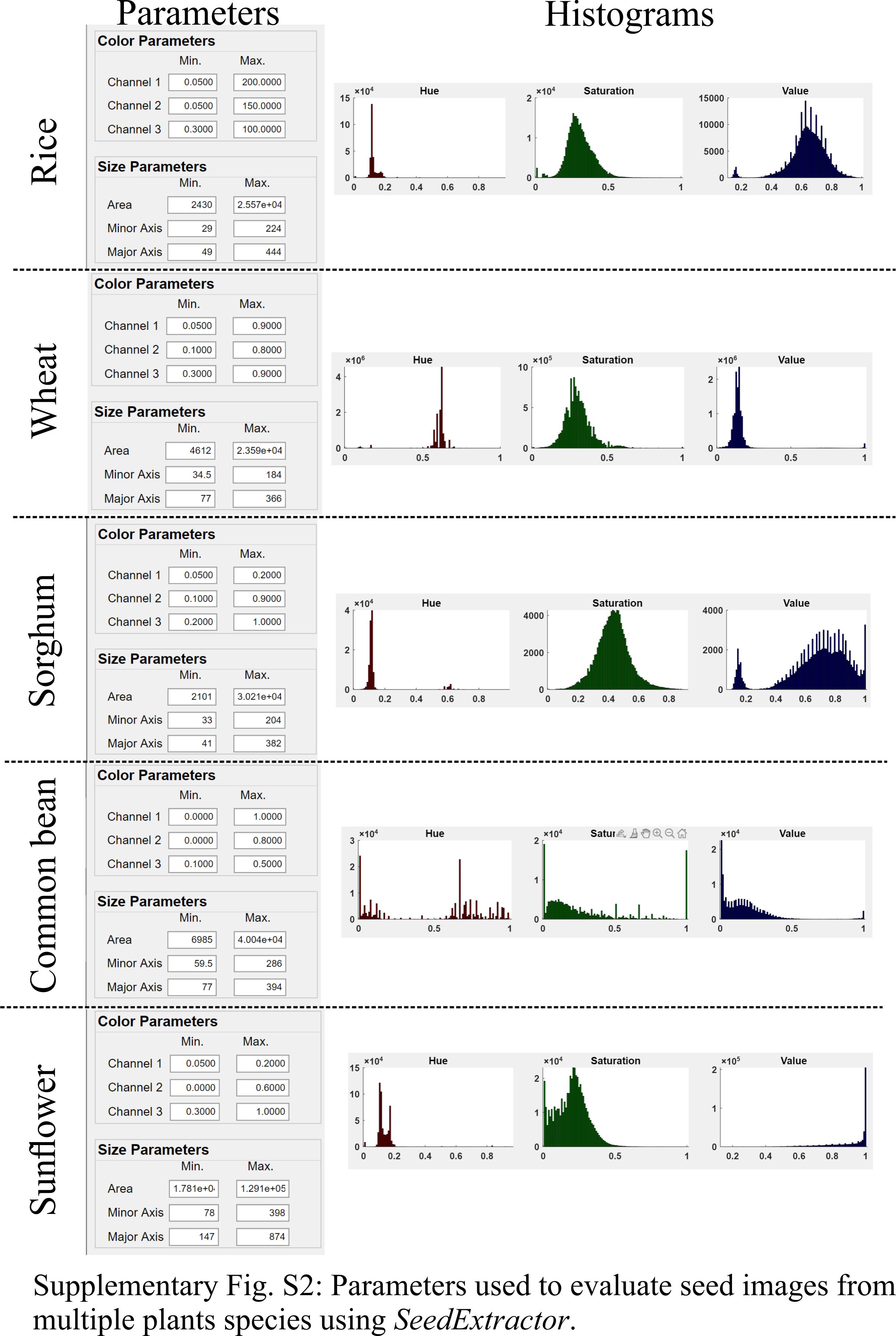

### Fig. S3

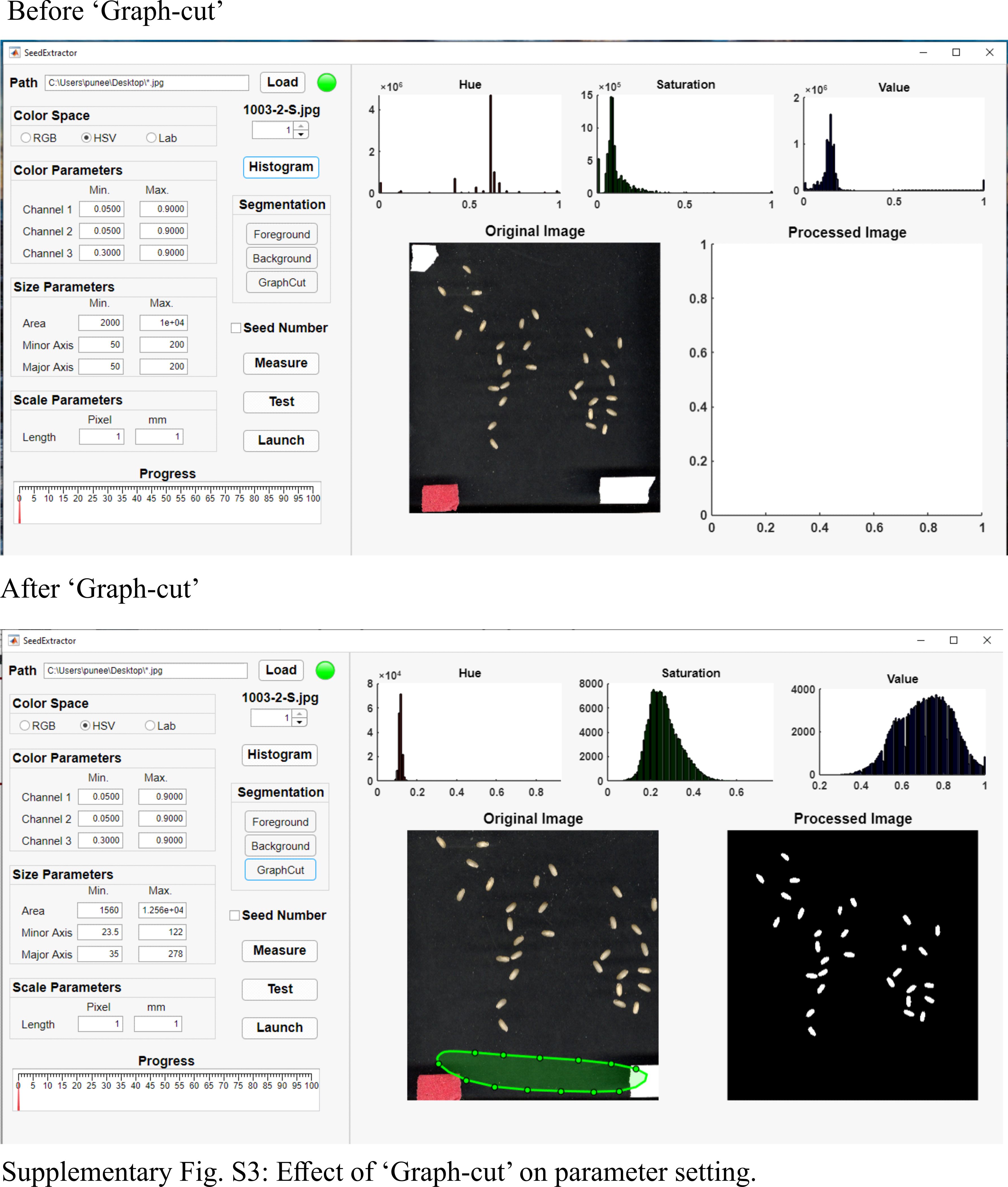

### Fig. S4

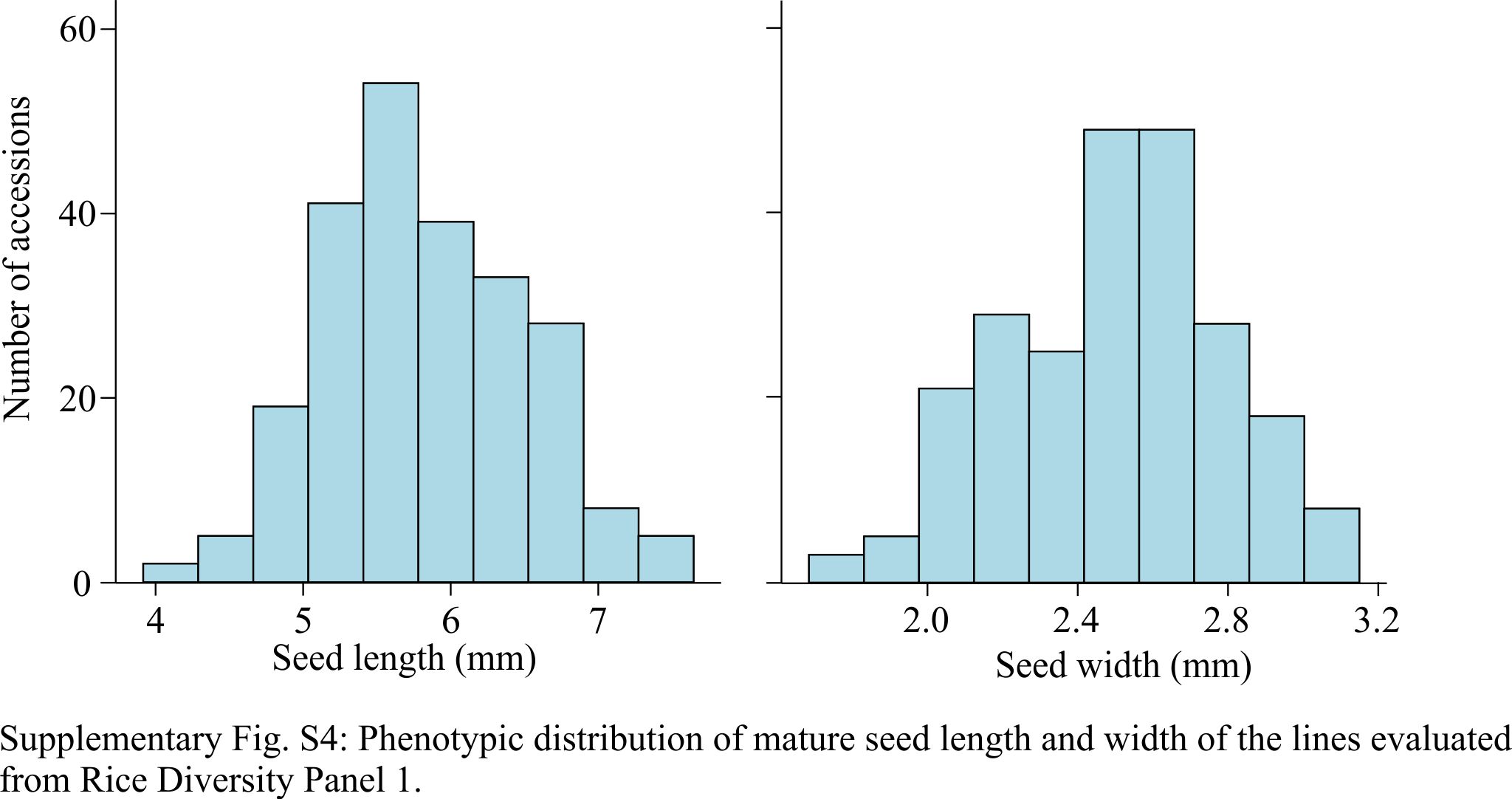
