## Supplementary material for "*SeedExtractor*: an open-source GUI for seed image analysis": SeedExtractor Guide Document

### Detailed step-by-step guide for using *SeedExtractor*

**Zhu and Paul *et al*., 2020**

#### Download and Launching the APPLICATION

1. Download MATLAB https://www.mathworks.com/products/matlab.html and *SeedExtractor* https://cse.unl.edu/~fzhu/SeedExtractor.zip.
2. Install the *SeedExtractor* application.
3. Open the MATLAB and go to APPS → My Apps → *SeedExtractor*.
4. Click on *SeedExtractor* to launch the application.

#### Loading the image and initiating the processing procedure

1. Once the application is launched, provide the complete **PATH** (FOLDER NAME***.jpg**) of the folder containing the scanned or camera-based images to be analyzed. * in the **PATH** represents that all the images in the respective folder need to be processed (Fig. 1: upper panel).
2. Click on the **LOAD** button. The light bulb adjacent to the **LOAD** button will change its colors (red to green) signifying that all image files are loaded.
3. One of the images from the folder will appear in **ORIGINAL IMAGE** space on the right panel of the application (Fig. 1: lower panel).
4. The user can use the **IMAGE INDEX** spinner to change the current image shown in the **ORIGINAL IMAGE** space to visualize other images in the parent folder.
5. Next, the user can pick the **COLOR SPACE** of choice: RGB, HSV or Lab.
6. The default color and size parameters are given by the application; however, the user can change or adjust the parameters as per their requirements.

   ***NOTE***: For color parameters, we highly recommend the user to consider the histogram generated after GraphCut as a guide to specify the minimum and maximum range for the specific channel. For seed size, the user needs to refer to the segmented or processed image and compare it to the original image to make appropriate changes in the seed size parameters for precise segmentation.

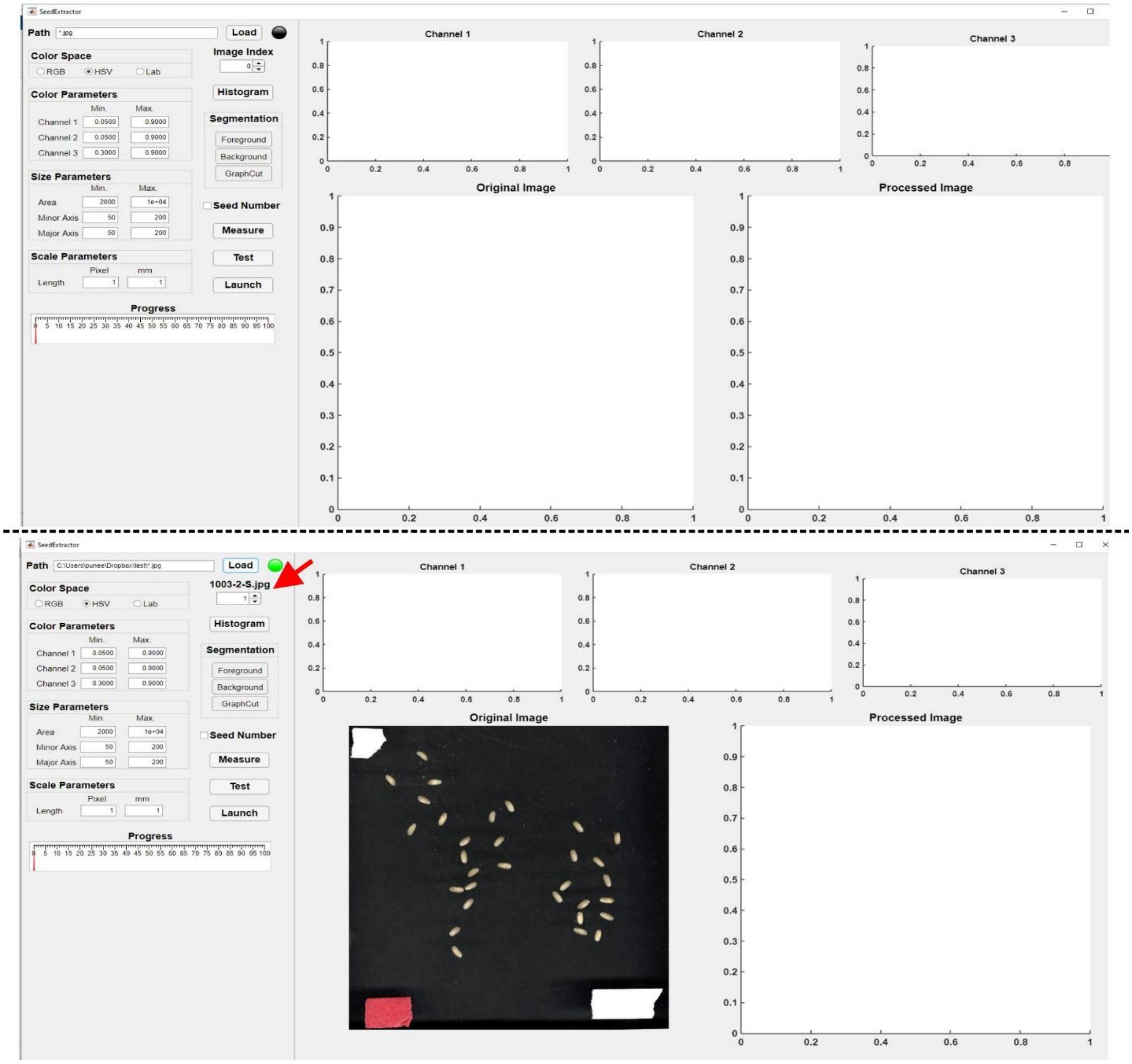


Fig. 1. The upper panel shows the graphical user interface of *SeedExtractor*. The lower panel shows a screenshot of the application showing the loaded image once the path of the folder containing the images is defined (FOLDER NAME*jpg). The light bulb next to the LOAD button will turn red during the process of image loading and will turn green once the loading is complete. The user can visualize other images in the parent folder using the spinner (designated by red arrow in the lower panel).

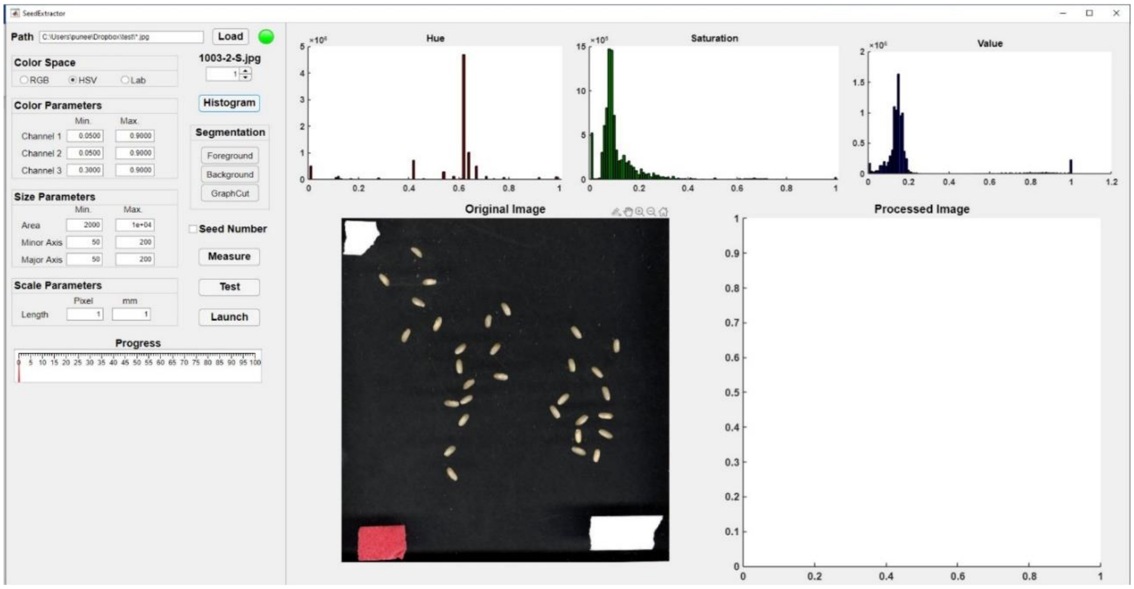


Fig. 2. If you click the HISTOGRAM button, the histograms will appear on the right section (upper panel) showing the distribution of colors in the three channels (Channel 1, 2, and 3 which in this case refer to Hue, Saturation, and Value) of the selected image.

#### Adjusting the parameters

Click on **HISTOGRAM** to visualize the ranges of the color distribution in the selected color space. **SEGMENTATION** function needs to be used to set the color and size parameters. For this, follow these steps:

1. If the seeds are relatively bigger, then the user can directly click on the **FOREGROUND** function and select the seed region on the **ORIGINAL IMAGE**. For selection, hold the left click on the mouse to draw on the region of interest, which will lead to appearance of small red circles on the selected seed region.
2. If the seeds are relatively smaller, then the user should move the cursor on the **ORIGINAL IMAGE** and a small **TASK BAR** will appear on the top of the image (Fig. 3A, shown with red arrow). Either use the wheel on your mouse or select the to **ZOOM IN** button (denoted by + symbol in a lens) to enlarge a specific region on the image that includes a single seed. If zoom in button is selected, the zoom in icon will turn blue (Fig. 3B).

   (*NOTE*: we recommend that the user to use the zoomed in step to define foreground, as it will increase the specificity of the segmentation procedure).
3. Select the area of interest, which will be the seed region (Fig. 3C, the application will make a blue box around the selected region). This will enlarge the region of interest (Fig. 3D).
4. Then, click on the **FOREGROUND** function for defining the seed region as foreground. For this, hold the left click on the mouse to draw on the region of interest, which will lead to appearance of

small red circles on the selected seed region (Fig. 3E). Next, the user should restore the normal view by clicking the **HOME** button on the task bar (Fig. 3F).

5. For selecting background of the image, click on **BACKGROUND** function and draw on the background of the image i.e. without the seed region (Fig. 3G). The background region under selection will be shown by green area.
6. Then, click on the **GRAPHCUT** to segment the foreground from the background. This step may take a few seconds and will generate a new histogram in each of the respective channel (Fig. 4). As a result, an image in the **PROCESSED IMAGE** space will appear (Fig. 4).
9. In case the processed image is not the accurate segmented version of the original image (if a seed region is missing for a seed or multiple seeds in the processed image compared to the original image), the user can *EITHER* try to re-select the foreground and background and reuse **GRAPHCUT** for segmentation *OR* manually re-set the color parameters, both minimum and maximum range, based on the distribution of histograms generated after using the **GRAPHCUT** for each of the channel. Similarly, the user can define the size parameters as well and set the minimum and maximum area, major and minor axis length for precise segmentation of the seed region.
10. This will help the user to ensure that the seeds in the original image are properly segmented in the processed image section. Nevertheless, the automatically set parameters offer reliable initial values for appropriate segmentation.

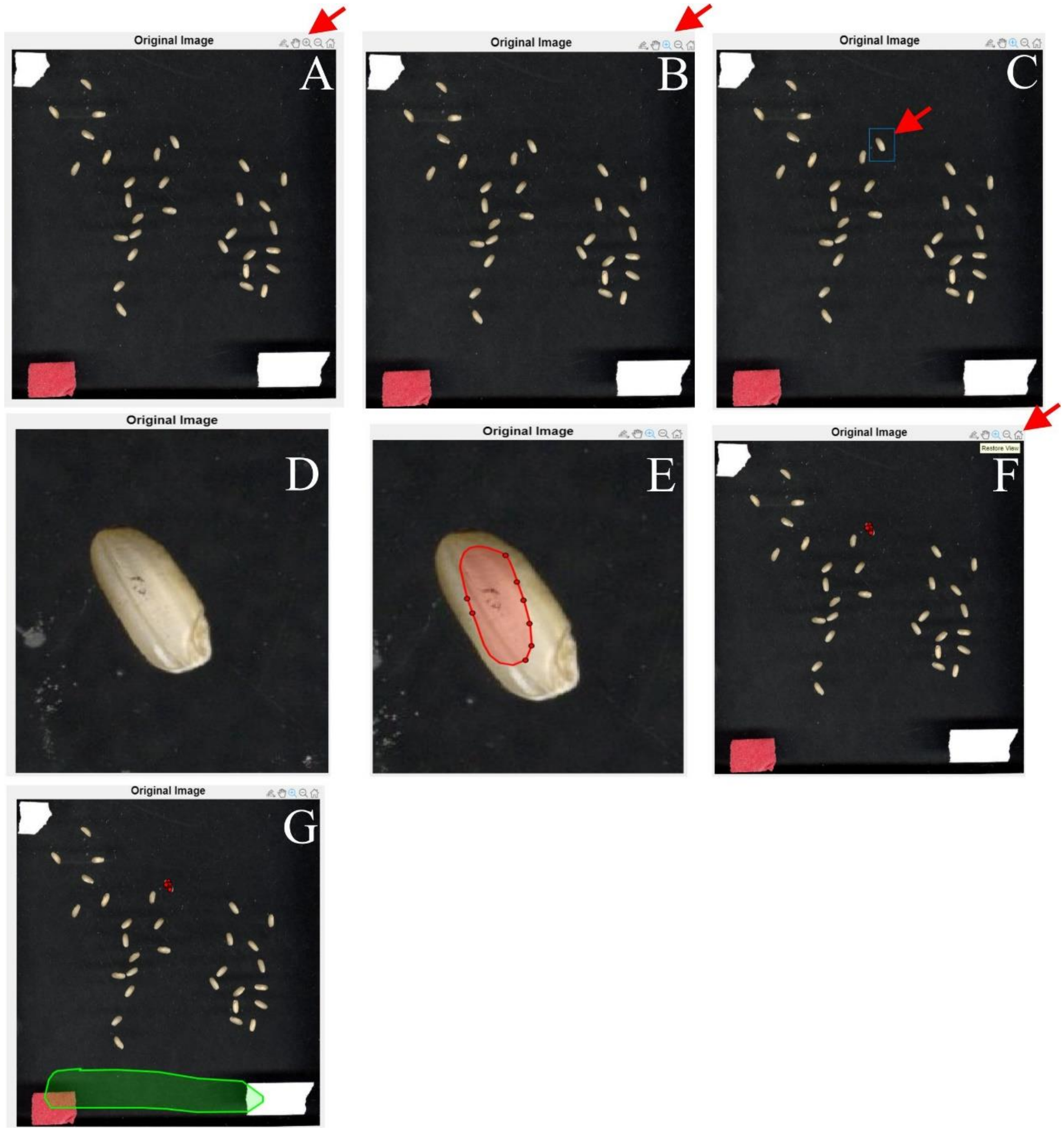


Fig. 3. Guide for selection of FOREGROUND and BACKGROUND. This step ensures appropriate and precise segmentation of the seed region using the GRAPHCUT procedure.

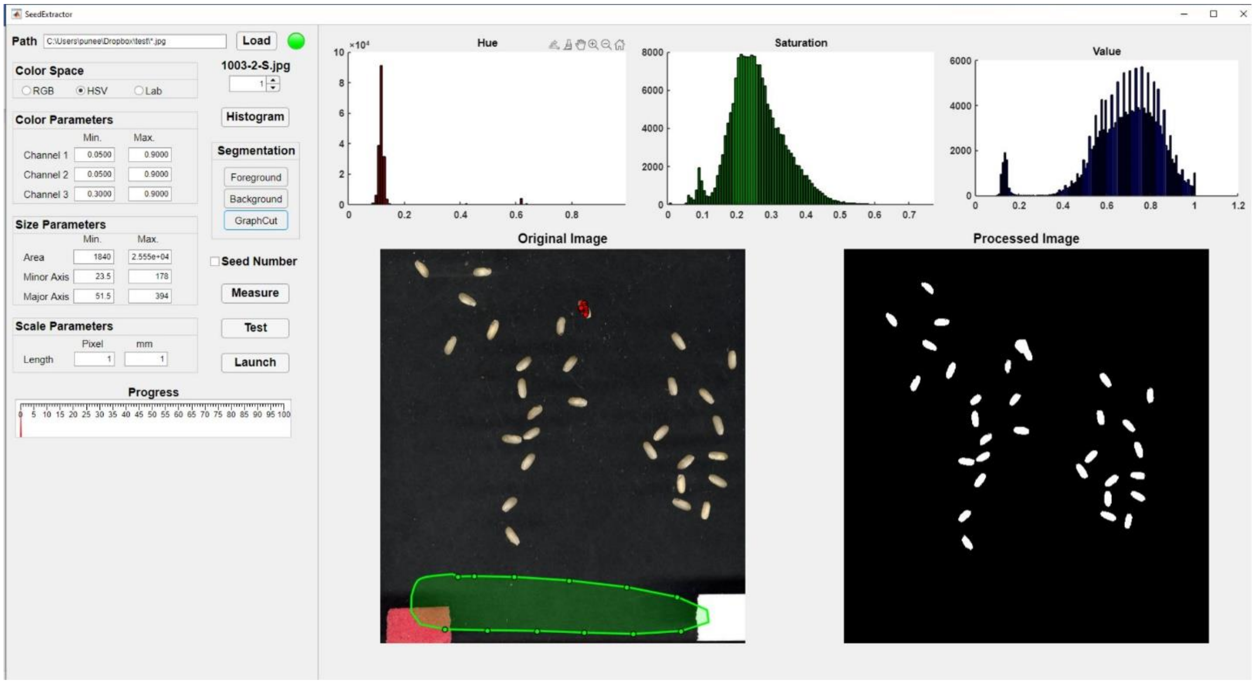


Fig. 4. Generation of the PROCESSED IMAGE and new histograms. The user can use the histograms (generated after using the GRAPHCUT for each of the channel) to manually re-set the color parameters (both minimum and maximum range for each of the channel). This will ensure optimal segmentation of the original image and generate a precise processed image.

#### Scale measurement

1. The application provides the measured seed parameters in pixels. However, the tool provides the flexibility for the user to measure objects that have been used as a scale in the image, which will transform the pixel length into millimeters (mm).
2. The user can again use the **ZOOM IN** function from the **TASK BAR** to select the scale region in the original image. Once the scale region is enlarged or zoomed, user can click on the **MEASURE** button to draw a straight blue line (Fig. 5, shown by red arrow). When the line is drawn, the pixel length of the blue line will appear in the 'Length (pixel)' textbox. The user can type the corresponding length of the blue line in the 'Length (mm)' textbox. Then, the application automatically converts those values into metric units in the output file.
3. The user can return to original image (normal view) by clicking the home icon on the **TASK BAR**.

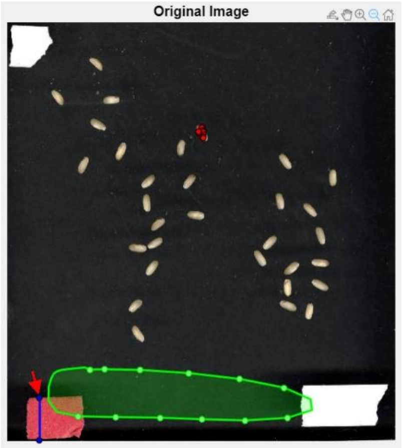


Fig. 5. Scale measurement. The drawing of straight blue line (signified by the red arrow) represents the scale (with the known measurements) defined by the user. By utilizing this function, the user can convert the scale from pixel length to metric length. For this, user can input the known or actual length of the blue line in the 'Length (mm)' textbox that corresponds to 'Length (pixel)' textbox. The application automatically converts those values into metric units in the output file.

#### Indexing the seed number and testing the parameters

1. The user can select the **SEED NUMBER** checkbox to specify individual seeds in the processed image.
2. The set parameters can be tested by clicking the **TEST** button. The seed numbers will be visible in small yellow boxes for each of the seed (Fig. 6).
3. User can look at the values generated by the tool in the MATLAB console.

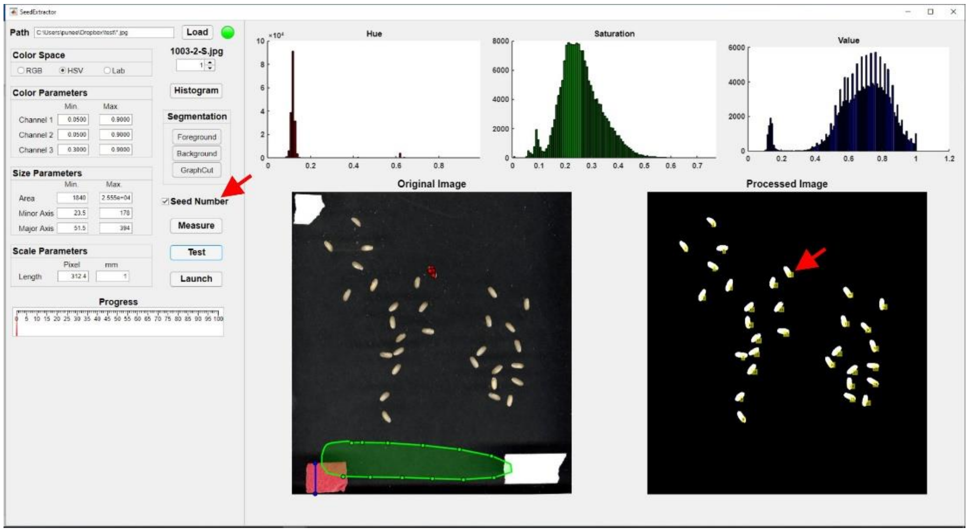


Fig. 6. Indexing the seeds. Clicking the SEED NUMBER function (shown by red arrow in the left panel), the user can define the seed number indices on the processed image and in the output file.

#### Batch processing

1. If the user decided on the parameters through testing, then **LAUNCH** button can clicked to initiate batch processing of the images. This will facilitate processing of all the images in the respective folder.
2. The progress of the processing can be monitored from the **PROGRESS** gauge.

#### Output files

1. For each processed image, the application will generate an output file in the parent folder. This file will contain trait information of each seed in that image.
2. Likewise, the mask of the seed regions from each image will be generated as a processed image in the parent folder. The indices of all the seed regions are marked in the processed image (if seed number checkbox is selected).
3. In addition, the user can download a combined file (*TotalResult.csv*) representing the average of a particular trait for all the seeds per image under the current path of MATLAB or from the MATLAB console (Fig. 7).

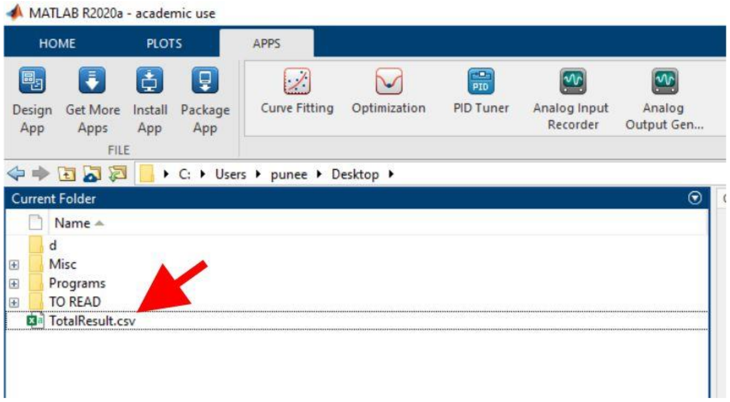


Fig. 7. The user can download combined file (*TotalResult.csv*) representing the average of a particular trait for all the seeds per image in the respective folder from the MATLAB console.
